# The Developing Human Connectome Project: typical and disrupted perinatal functional connectivity

**DOI:** 10.1101/2020.01.20.912881

**Authors:** Michael Eyre, Sean P Fitzgibbon, Judit Ciarrusta, Lucilio Cordero-Grande, Anthony N Price, Tanya Poppe, Andreas Schuh, Emer Hughes, Camilla O’Keeffe, Jakki Brandon, Daniel Cromb, Katy Vecchiato, Jesper Andersson, Eugene P Duff, Serena J Counsell, Stephen M Smith, Daniel Rueckert, Joseph V Hajnal, Tomoki Arichi, Jonathan O’Muircheartaigh, Dafnis Batalle, A David Edwards

## Abstract

The Developing Human Connectome Project (dHCP) is an Open Science project which provides the first large sample of neonatal functional MRI (fMRI) data with high temporal and spatial resolution. This data enables mapping of intrinsic functional connectivity between spatially distributed brain regions under normal and adverse perinatal circumstances, offering a framework to study the ontogeny of large-scale brain organisation in humans. Here, we characterise in unprecedented detail the maturation and integrity of resting-state networks (RSNs) at normal term age in 337 infants (including 65 born preterm).

First, we applied group independent component analysis (ICA) to define 11 RSNs in term-born infants scanned at 43.5-44.5 weeks postmenstrual age (PMA). Adult-like topography was observed in RSNs encompassing primary sensorimotor, visual and auditory cortices. Among six higher-order, association RSNs, analogues of the adult networks for language and ocular control were identified, but a complete default mode network precursor was not. Next, we regressed the subject-level datasets from an independent cohort of infants scanned at 37-43.5 weeks PMA against the group-level RSNs to test for the effects of age, sex and preterm birth. Brain mapping in term-born infants revealed areas of positive association with age across four of six association RSNs, indicating active maturation in functional connectivity from 37 to 43.5 weeks PMA. Female infants showed increased connectivity in inferotemporal regions of the visual association network. Preterm birth was associated with striking impairments of functional connectivity across all RSNs in a dose-dependent manner; conversely, connectivity of the superior parietal lobules within the lateral motor network was abnormally increased in preterm infants, suggesting a possible mechanism for specific difficulties such as developmental coordination disorder which occur frequently in preterm children.

Overall, we find a robust, modular, symmetrical functional brain organisation at normal term age. A complete set of adult-equivalent primary RSNs is already instated, alongside emerging connectivity in immature association RSNs, consistent with a primary-to-higher-order ontogenetic sequence of brain development. The early developmental disruption imposed by preterm birth is associated with extensive alterations in functional connectivity.

## Introduction

The Developing Human Connectome Project (dHCP) is an Open Science project funded by the European Research Council to provide a large dataset of functional and structural brain images from 20 to 44 weeks of gestational age (GA). This enables the characterisation of 4-dimensional (three spatial dimensions and time) connectivity maps, which map the trajectories of human brain development to improve understanding of normal development and allow earlier detection and intervention for neurological and psychological disorders.

This paper analyses functional connectivity at the time of normal birth in infants born at term and preterm. Temporal coherences in the blood-oxygen–level-dependent (BOLD) contrast measured with resting-state functional magnetic resonance images (rs-fMRI) can be spatiotemporally decomposed into resting-state networks (RSNs) (Damoiseaux et al., 2006) (Biswal et al., 1995), predominantly at low frequency (< 0.1 Hz) (Cordes et al., 2001), distinct from cardiovascular signal (De Luca et al., 2006). Whilst RSNs have been extensively and robustly characterized in the mature brain, previous studies of RSN development in newborn infants have been limited by smaller sample sizes. The dHCP provides the first high-quality, large-scale, 4-dimensional dataset of structural-functional connectivity at this critical period of development, enabling us to address two key questions. Firstly, are higher-order RSNs such as the default-mode network (DMN) (Raichle et al., 2001) instated with adult topology in the neonatal period? Some find analogues of these at term-equivalent age (TEA) (Doria et al., 2010; Fransson et al., 2009; Fransson et al., 2007; Smyser et al., 2010; Smyser et al., 2016) while others locate their origin in later infancy or early childhood, contemporaneous with the emergence of the higher cognitive abilities these networks are believed to support (Gao et al., 2014; Gao et al., 2015). Secondly, what is the effect of preterm birth on RSN development? Although various alterations in the complexity, scope, strength and efficiency of functional connectivity in preterm-at-term infants have been reported (Ball et al., 2016; Bouyssi-Kobar et al., 2019; Doria et al., 2010; Smyser et al., 2010; Smyser et al., 2016; Toulmin et al., 2015), the majority of studies lack the large numbers of control subjects required to characterise these effects with precision.

The mature adult RSNs are well characterised, with high intra-subject reproducibility (Finn et al., 2015; Wang et al., 2015), largely consistent topology across healthy subjects, and anatomical mapping that reinforces both structural and task-fMRI-derived parcellations of the cortex (Glasser et al., 2016). The identification of fMRI-RSN signatures associated with disease offers considerable translational potential due to rs-fMRI’s relatively straightforward and widely-used acquisition, whole-brain coverage, and high spatial resolution compared to other functional imaging methods. The achievement of this in the immature brain requires a complete account of RSN ontogeny. The developing CNS shows spontaneous, patterned, correlated intrinsic activity from early prenatal life (reviewed in (Blankenship and Feller, 2010; Keunen et al., 2017; Vasung et al., 2019); immature RSNs can be identified from as early as 26 weeks postmenstrual age (PMA) in the preterm infant (Doria et al., 2010; Smyser et al., 2010). By TEA, RSNs bearing closer resemblance to adult RSNs are observed in both term- and preterm-born infants (Doria et al., 2010; Fransson et al., 2011; Fransson et al., 2009; Fransson et al., 2007; Gao et al., 2014; Gao et al., 2015; Smyser et al., 2016). However, TEA encompasses a critical period of brain development in which there is intense myelination of white matter (reviewed in (Dubois et al., 2014) and rapid expansion in both the size and gyrification of the cerebral cortex (Dubois et al., 2019; Shimony et al., 2016). Dense sampling across the age range is therefore required to map the associated changes in functional connectivity.

Here we apply a data-driven approach to 337 rs-fMRI datasets acquired in term and preterm infants between 37 and 44.5 weeks PMA. We first defined a normative set of RSNs in a sub-sample of term-born infants scanned at 43.5-44.5 weeks PMA using probabilistic independent component analysis (ICA) (Beckmann and Smith, 2004). ICA is a dimensionality reduction technique which decomposes data into a set of components with maximal statistical independence; applied to rs-fMRI, ICA can reveal large-scale brain networks without requirement for a predefined model of network structure. We then regressed subject-level data from term and preterm infants scanned at 37-43.5 weeks PMA against these networks. The resulting whole-brain correlation maps enabled us to both characterise the ontogeny of individual RSNs, and investigate the influence of prematurity on cortical functional connectivity.

## Materials and methods

### Subjects

Research participants were prospectively recruited as part of the dHCP, an observational, cross-sectional Open Science programme approved by the UK National Research Ethics Authority (14/LO/1169). Written consent was obtained from all participating families prior to imaging. We selected 416 structural-functional datasets acquired at TEA from the 2019 (second) dHCP data release. Only infants scanned at 37-44.5 weeks PMA in term-born infants, or 37-43.5 weeks PMA in preterm-born infants, were considered for inclusion. One infant was included twice due to two datasets being acquired at different ages; only the second dataset was used. Thirty-five infants were excluded due to a history of neurodevelopmental disorder in a first-degree relative. Forty-three were excluded due to motion (see **Functional data pre-processing**). The final study population therefore consisted of 337 infants, divided into three groups: term-born infants scanned at 43.5-44.5 weeks PMA, who were used to define the normative set of RSNs and excluded from all subsequent subject-level analyses (i); and the remaining infants scanned at 37-43.5 weeks PMA, including both term-born (ii) and preterm-born (iii) infants (**Table 1**).

### MR data acquisition

Neuroimaging was acquired in a single scan session for each infant at the Evelina Newborn Imaging Centre, Evelina London Children’s Hospital, using a 3-Tesla Philips Achieva system (Philips Medical Systems, Best, The Netherlands). All infants were scanned without sedation in a scanner environment optimized for safe and comfortable neonatal imaging, including a dedicated transport system, positioning device and a customized 32-channel receive coil, with a custom-made acoustic hood (Hughes et al., 2017). MR-compatible ear putty and earmuffs were used to provide additional acoustic noise attenuation. Infants were fed, swaddled and comfortably positioned in a vacuum jacket prior to scanning to promote natural sleep. All scans were supervised by a neonatal nurse and/or paediatrician who monitored heart rate, oxygen saturation and temperature throughout the scan.

High-temporal-resolution BOLD fMRI optimized for neonates was acquired over 15 minutes 3 seconds (2300 volumes) using a multislice gradient-echo echo planar imaging (EPI) sequence with multiband excitation (multiband factor 9). Repetition time (TR) was 392 milliseconds, echo time (TE) was 38 milliseconds, flip angle was 34°, and the acquired spatial resolution was 2.15 mm isotropic (Price et al., 2015). For registration of the fMRI data, high-resolution T1- and T2-weighted anatomical imaging was also acquired in the same scan session, with a spatial resolution of 0.8 mm isotropic (T1w: field of view 145 × 122 × 100 mm, TR 4795 ms; T2w: field of view 145 × 145 × 108 mm, TR 12000 ms, TE 156 ms).

### Functional data pre-processing

Data were pre-processed using an in-house pipeline optimized for neonatal imaging and specifically developed for the dHCP, detailed in Fitzgibbon et al. (2019). In brief, susceptibility dynamic distortion together with intra- and inter-volume motion effects were corrected in each subject using a bespoke pipeline including slice-to-volume and rigid-body registration (Andersson et al., 2018; Andersson et al., 2017; Andersson et al., 2001; Andersson et al., 2003). In order to regress out signal artifacts related to head motion, cardiorespiratory fluctuations and multiband acquisition (Salimi-Khorshidi et al., 2014), 24 extended rigid-body motion parameters were regressed together with single-subject ICA noise components identified with the FSL FIX tool (Oxford Centre for Functional Magnetic Resonance Imaging of the Brain’s Software Library, version 5.0). Denoised data were registered into T2w native space using boundary-based registration (Greve and Fischl, 2009) and non-linearly registered to a standard space based in a weekly template from the dHCP volumetric atlas (Schuh et al., 2018) using a diffeomorphic multimodal (T1/T2) registration (Avants et al., 2008).

While the fMRI preprocessing pipeline for the dHCP (Fitzgibbon et al., 2019) addresses the potential problem of head motion in rs-fMRI data (Power et al., 2012; Satterthwaite et al., 2012), motion is also a surrogate marker of the arousal state of the infant, which interacts with the underlying neural activity (Denisova, 2019; Whitehead et al., 2018). To address this issue, we opted for a conservative approach consisting in the selection of a continuous sub-sample of the data (~70%) with lowest motion for each subject, and excluding those subjects with a high level of motion from further analyses. Specifically, volumes with DVARS (the root mean square intensity difference between successive volumes) higher than 1.5 IQR above the 75^th^ centile, after motion and distortion correction, were considered as motion outliers (Fitzgibbon et al., 2019); within each acquired dataset (2300 volumes), the continuous set of 1600 volumes with the minimum number of motion-outlier volumes was identified, and the dataset cropped accordingly for all subsequent analyses. Subjects with more than 160 motion-outlier volumes (10% of the cropped dataset) were excluded entirely. This allowed us to minimise the potential effect of different states of arousal even after appropriately denoising the data. The number of motion-outlier volumes remaining in the cropped dataset was recorded for each subject and included as a covariate in all subsequent regression analyses. The median number of motion-outlier volumes in the term-born group was 49.5 (IQR 27-86.5) and in the preterm-born group was 34 (IQR 12-83) (p = 0.052, Mann-Whitney U test).

### Functional data analysis

#### Group-level network definition

We first defined the normative set of RSNs by group ICA in 24 healthy term-born infants scanned at 43.5-44.5 weeks PMA. These subjects were excluded from all subsequent regression analyses. Probabilistic group ICA by temporal concatenation across subjects was carried out using FSL MELODIC (Beckmann and Smith, 2004). The ICA dimensionality was set at 30, representing a pragmatic balance between robustness and interpretability (as in (Toulmin et al., 2015)). The output comprised 30 group-average spatial maps representing 30 independent components. The maps were visually inspected and each component manually labelled as RSN or noise, following guidelines in Fitzgibbon et al. (2019).

#### Subject-level analyses

We next regressed the group-level spatial maps into the subject-level 4D space-time datasets of the subjects scanned at 37-43.5 weeks PMA (248 term-born, 65 preterm-born). Specifically, the group-level spatial maps (including both RSN signal and artifact components) were used to generate subject-specific versions of the spatial maps and associated time series using dual regression (Nickerson et al., 2017). First, for each subject, the set of group-level RSN spatial maps was regressed (as spatial regressors in a multiple regression) into the subject’s 4D space-time dataset. This resulted in a set of subject-specific time series, one per group-level spatial map. Next, those timeseries were regressed (as temporal regressors, again in a multiple regression) into the same 4D dataset, resulting in a set of subject-specific spatial maps, one per group-level spatial map.

We then performed cross-subject analysis using general linear models (GLM) to test for the effects of group (term vs. preterm birth, sex) and continuous variables (GA at birth, PMA at scan) on the subject-level spatial maps, including the number of motion-compromised volumes as a nuisance covariate. A further group-level analysis was conducted in which term-born infants were separated into weekly bins according to their PMA at scan, enabling group-average maps of functional connectivity at each week of brain development to be generated. For this we entered data from the 20 subjects in each bin with the lowest number of postnatal days of life at time of scan, to maximise similarity between groups for meaningful visual comparison.

To further quantify longitudinal changes in within-network functional connectivity, we analysed the relationship between PMA at scan and a derived parameter we term ‘core network strength’. This measure was determined for each RSN for each subject by masking the RSN-specific spatial map (the output of stage two of dual regression) by the corresponding group-ICA network template thresholded at *Z* > 3, then calculating the mean β parameter value (regression coefficient) within the masked image. The partial Spearman’s correlation between core network strength and PMA at scan was calculated in term-born infants while controlling for sex and motion, and a GLM was used to test for group differences in core network strength between term and preterm infants while controlling for PMA at scan, sex and motion.

#### Statistical tests

Voxelwise statistical tests were implemented in FSL *randomise* (Winkler et al., 2014) using threshold-free cluster enhancement (Smith and Nichols, 2009) with 5000 permutations. As all contrasts were two-tailed, family-wise error-rate (FWE) corrected (for multiple comparisons across voxels) p-values less than 0.025 were accepted as significant. Correlation and GLM analyses of core network strength were implemented in Python 3.7 with *pingouin* 0.2.9 and *statsmodels.api* 0.10.1.

#### Anatomical localisation and data visualisation

Results were localised in the standard space using an in-house adaptation of the neonatal version (Shi et al., 2011) of the AAL atlas (Tzourio-Mazoyer et al., 2002), projected to the 40-week high-resolution neonatal dHCP template (Schuh et al., 2018).

Data were displayed using *FSLeyes* for planar visualisation and Connectome Workbench for cortical surface visualisation.

##### Data availability

The dHCP is an open-access project. The imaging and collateral data used in this study were included in the 2019 (second) dHCP data release, which can be downloaded by registering at https://data.developingconnectome.org/

## Results

### Resting-state networks

Eleven RSNs were identified by group ICA in a sub-sample of term-born infants scanned between 43.5 and 44.5 weeks PMA (*n* =24), who were excluded from any further analyses. Five RSNs included primary motor or sensory cortical areas and were categorised as primary networks (**Fig. 1A**): medial motor, lateral motor, somatosensory, auditory and visual. The remaining six were categorised as association networks (**Fig. 1C**): motor association (including the premotor and supplementary motor areas), temporoparietal (including Broca’s area and the extended Wernicke’s area), posterior parietal (including the precuneus and posterior cingulate cortices), frontoparietal (including the frontal, supplementary and parietal eye fields), prefrontal and visual association. The full cortical surface parcellation is provided in **Supplementary Fig. S1** and the **Supplementary Video**.

**Figure 1.**
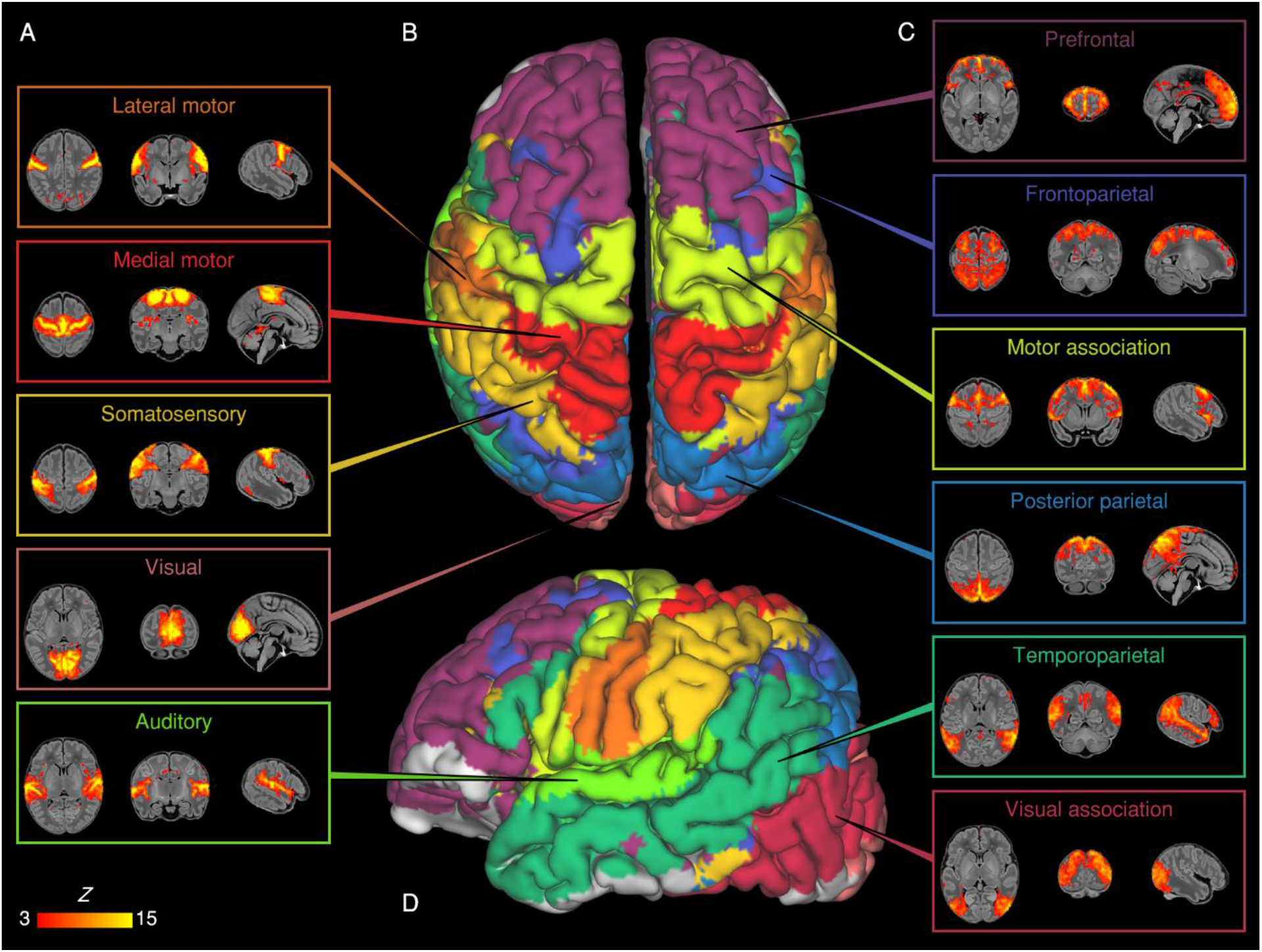
Resting-state networks identified by group independent component analysis. Spontaneous BOLD activity patterns (RSNs) derived from group ICA in 24 term-born infants scanned at 43.5-44.5 weeks PMA. Panels: Example axial, coronal, and sagittal slices for meaningful spatial patterns in primary (A) and association (C) RSNs, thresholded at Z > 3 and overlaid on a T1 structural template, displayed in radiological convention. Centre: Functional parcellation of the brain using a ‘winner-takes-all’ approach based on the RSNs from group ICA. RSNs were spatially smoothed and thresholded at Z > 1 prior to determination of the ‘winning’ RSN at each voxel. The resulting volume was projected to the midthickness cortical surface using enclosed (nearest neighbour) volume-to-surface mapping, here displayed on the pial surface of an individual subject scanned at 42 weeks PMA and viewed from the dorsal (B) and left lateral (D) aspects.

### Effect of postmenstrual age at scan

To characterise normal maturation in functional connectivity from 37-43.5 weeks in term-born infants, we analysed the association between previously calculated RSNs independently regressed to each subject and PMA at scan, while controlling for sex and motion. Brain regions showing increased connectivity with older PMA at scan were identified in four RSNs, all association networks (**Fig. 2**). There were no brain tissue regions showing negative association.

**Figure 2.**
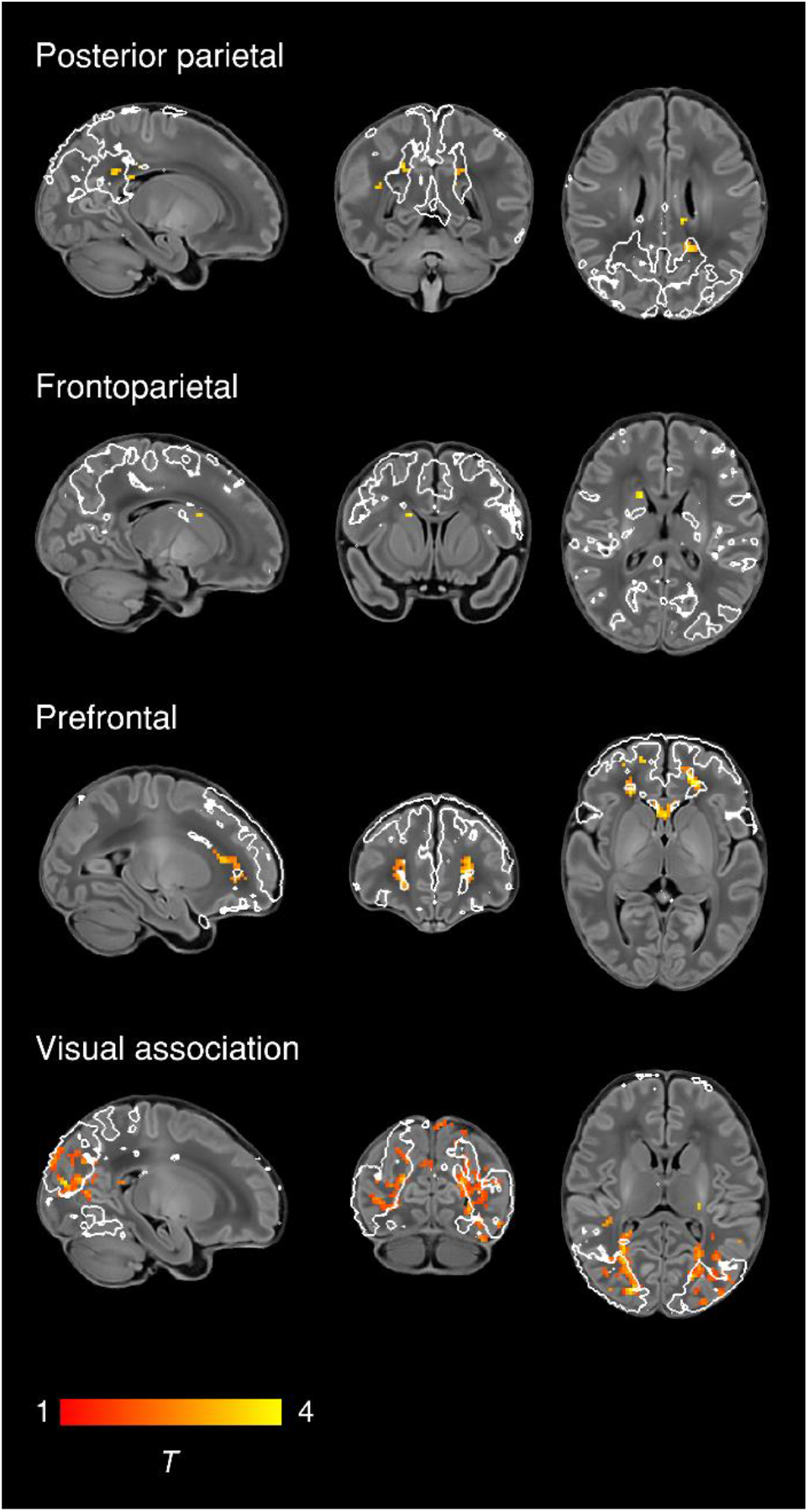
Effect of postmenstrual age at scan on functional connectivity. Brain regions showing increased functional connectivity with older PMA at scan in term-born infants scanned at 37-43.5 weeks PMA. Example sagittal, coronal, and axial slices for meaningful spatial patterns in four RSNs are shown, overlaid on a T1 structural template and displayed in radiological convention. T-statistic maps were thresholded at p < 0.025 (FWE corrected). White lines represent the outlines of the group-ICA RSNs, thresholded at Z > 3.

To further illustrate maturational changes in functional connectivity, we produced spatial maps of average network structure in term-born infants categorised into weekly groups according to their PMA at scan, while controlling for sex and motion (**Fig. 3**).

**Figure 3.**
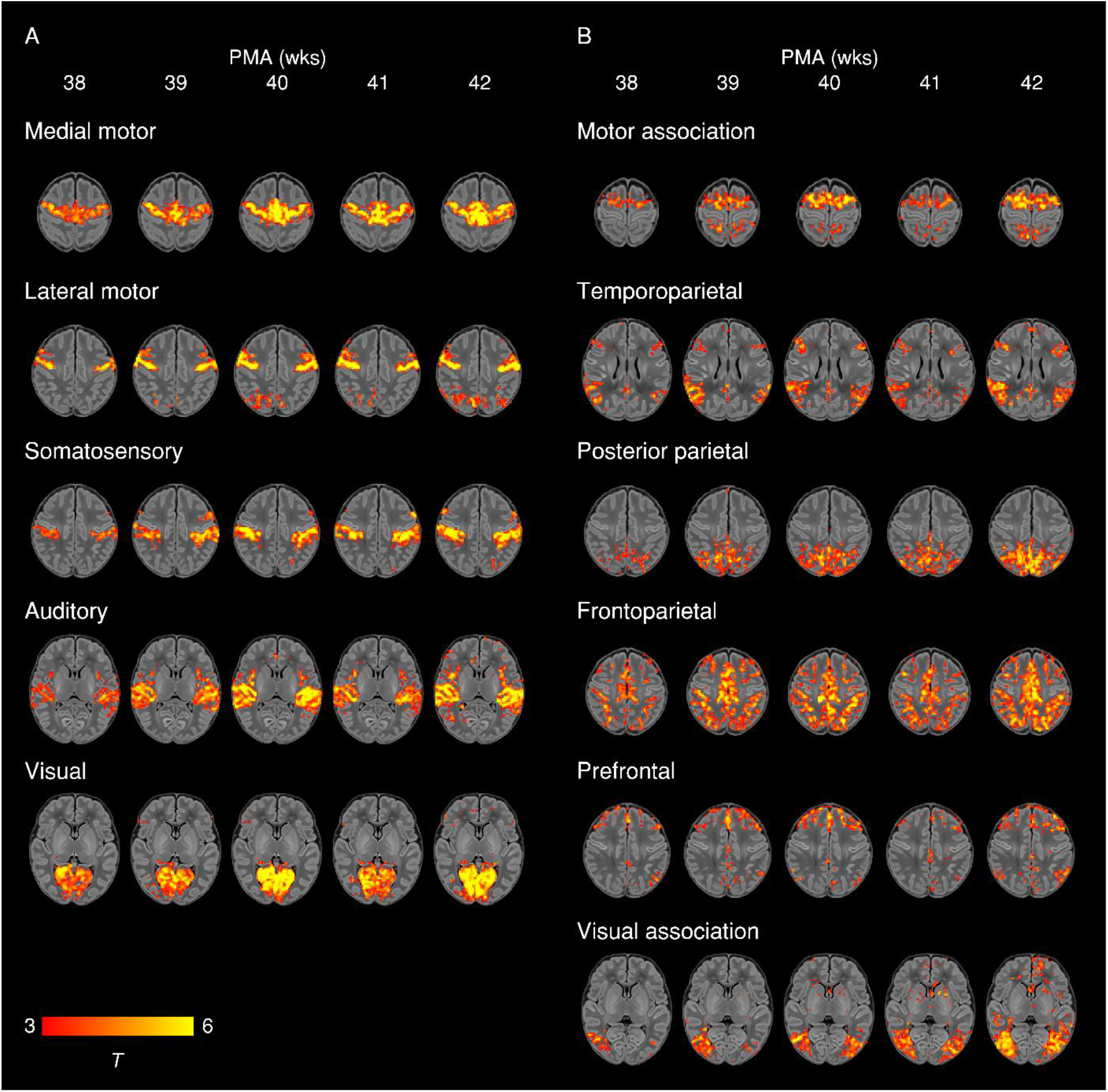
Weekly maturation in functional network structure at term-equivalent age. Group-average t-statistic maps of functional connectivity in term-born infants scanned at 37.5-42.5 weeks PMA, grouped into weekly bins by PMA at scan. Within each bin 20 subjects with the lowest postnatal age at time of scan were selected. Example axial slices for meaningful spatial patterns in primary (A) and association (B) RSNs are shown, overlaid on a T1 structural template and displayed in radiological convention. Results were thresholded at p < 0.05 (FWE corrected).

To further quantify longitudinal changes in within-network functional connectivity, we analysed the relationship between PMA at scan and a derived parameter we term ‘core network strength’, defined as the mean β parameter value in each subject’s RSN-specific spatial map (the outputs of stage two of dual regression) after masking by the corresponding group-ICA network template thresholded at *Z* > 3. Three RSNs showed a positive partial correlation between PMA at scan and core network strength (**Fig. 4**). There were no RSNs with negative correlation.

**Figure 4.**
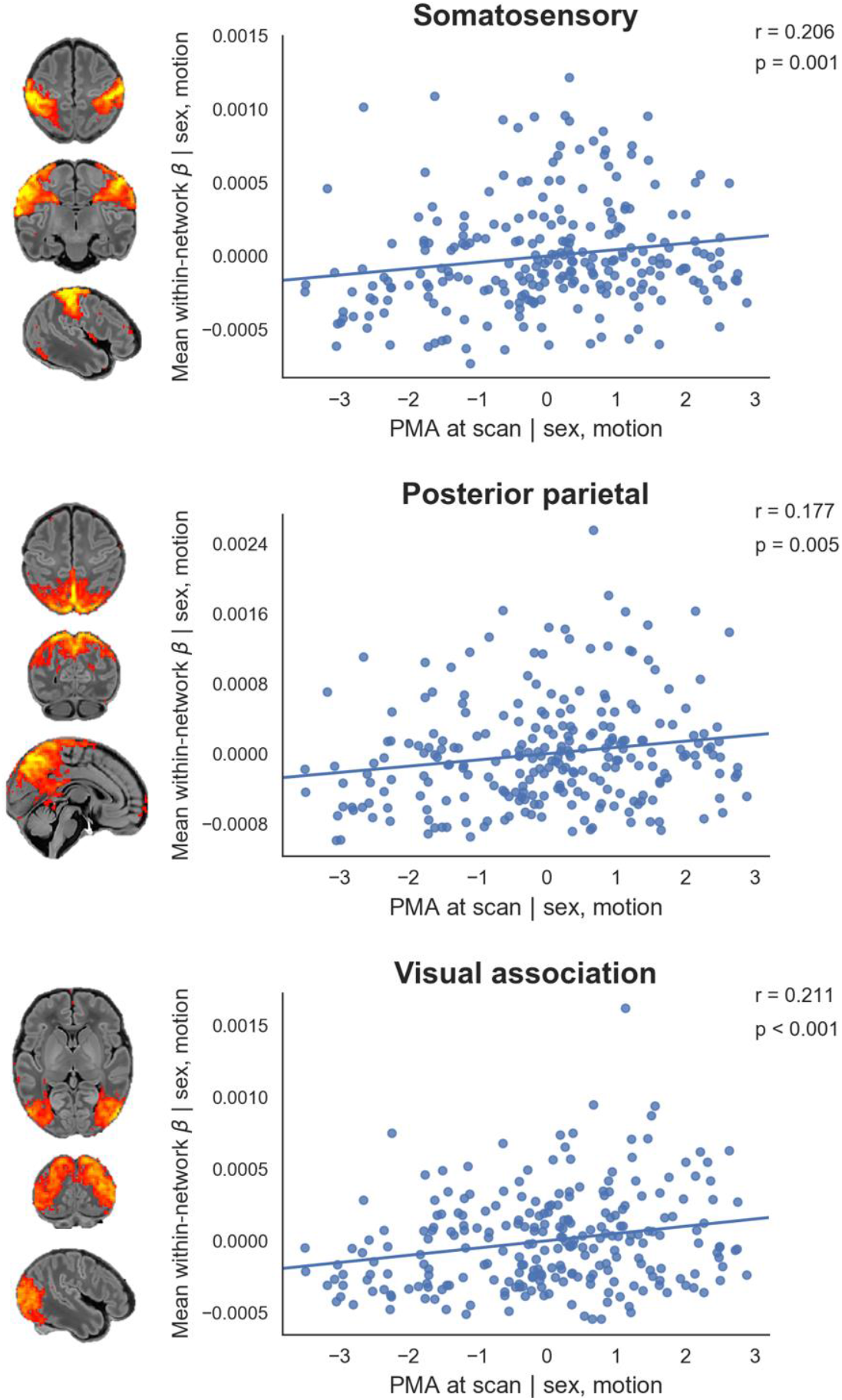
Relationship between postmenstrual age at scan and core network strength. Relationship between the residuals (after correcting for sex and motion) for PMA at scan and core network strength in term-born infants scanned at 37-43.5 weeks PMA. Core network strength was defined as the mean β parameter value in each subject’s RSN-specific spatial map after masking by the corresponding group-ICA network template thresholded at Z > 3. Partial Spearman’s correlation coefficients and associated p values are displayed for the three RSNs significant at p < 0.025. Example axial, coronal and sagittal slices for meaningful spatial patterns in the corresponding group-ICA network templates are shown for reference.

### Effect of Sex

To determine differences in functional connectivity between male and female infants we analysed this as a group effect, while controlling for GA at birth, PMA at scan and motion. Female infants showed increased connectivity of inferior occipito-temporal regions (including the posterior fusiform gyrus) within the visual association network (**Fig. 5**).

**Figure 5.**
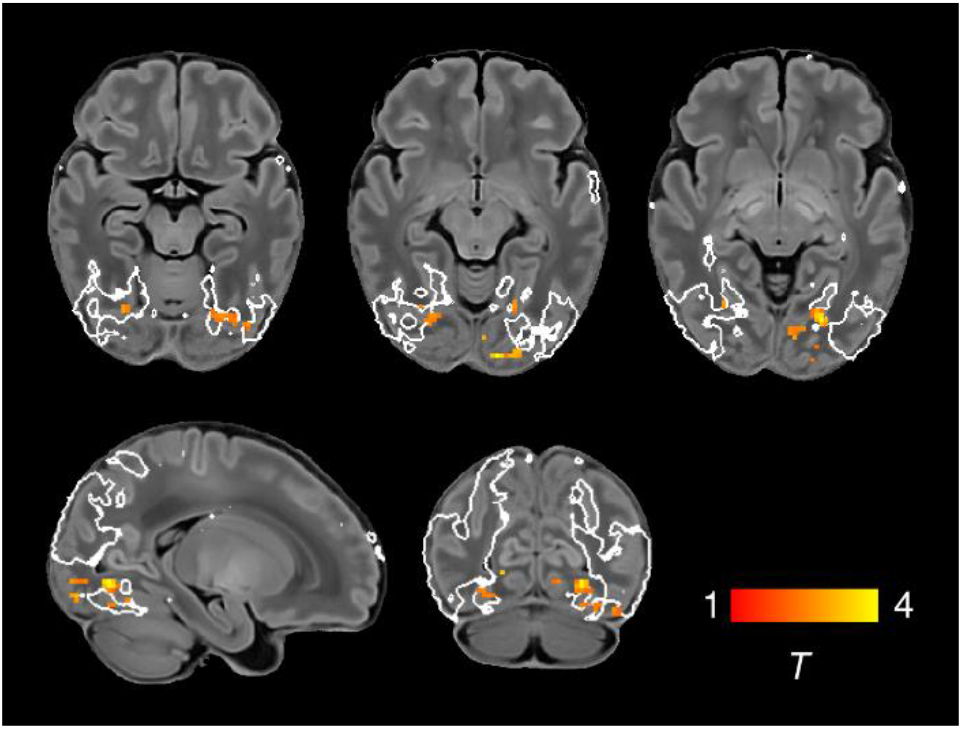
Increased functional connectivity in the visual association network in female infants. Brain regions showing increased functional connectivity within the visual association RSN in female infants. Example axial, sagittal and coronal slices for meaningful spatial patterns are shown, overlaid on a T1 structural template and displayed in radiological convention. T-statistic maps were thresholded at p < 0.025 (FWE corrected). White lines represent the outline of the group-ICA visual association network, thresholded at Z > 3.

### Effect of preterm birth

To determine differences in functional connectivity between term- and preterm-born infants we first analysed this as a group effect, while controlling for PMA at scan, sex and motion. There was extensive impairment of functional connectivity across all RSNs in preterm-born infants; uncorrected core network strength was 23-41% reduced relative to term-born infants across the 11 networks (all *p* < 0.001, independent samples t-tests) (**Fig. 6**). Conversely, preterm-born infants showed increased connectivity of the bilateral superior parietal lobule within the lateral motor network (**Fig. 6**). The association of younger GA at birth with impaired functional connectivity was replicated across all networks in a separate analysis in which GA at birth was entered as a continuous variable, indicating a dose-dependent effect of prematurity on functional connectivity (**Supplementary Fig. S2**).

**Figure 6.**
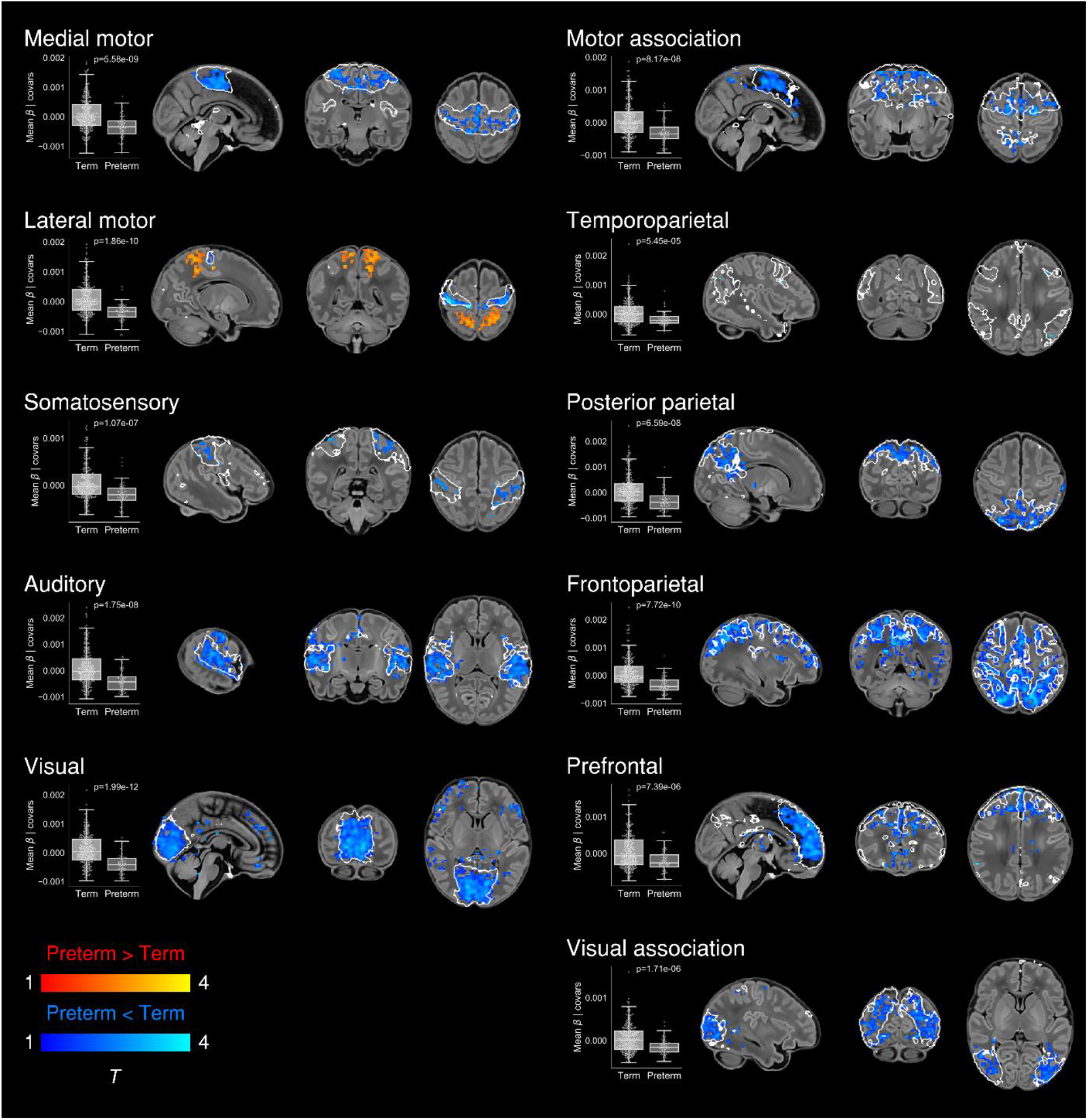
Effect of preterm birth on functional connectivity. Group differences in functional connectivity between term- and preterm-born infants scanned at 37-43.5 weeks PMA. Coloured t-statistic maps thresholded at p < 0.025 (FWE corrected) show brain regions with reduced (blue) or increased (red-yellow) connectivity in preterm-born infants. Example sagittal, coronal, and axial slices for meaningful spatial patterns within each RSN are shown, overlaid on a T1 structural template and displayed in radiological convention. White lines represent the outlines of the group-ICA RSNs, thresholded at Z > 3. Boxplots show group differences in core network strength after regressing out PMA at scan, sex and motion. Core network strength was defined as the mean β parameter value in each subject’s RSN-specific spatial map after masking by the corresponding group-ICA network template thresholded at Z > 3. P-values relate to the term vs. preterm group contrast in a GLM in which core network strength was the dependent variable and PMA at scan, sex and motion were controlled for as nuisance covariates.

## Discussion

In this large cohort of newborn infants we provide detailed characterization of the maturational trajectories of normal functional network development at TEA, and show that the early developmental disruption imposed by preterm birth is associated with significant and widespread alterations in functional connectivity.

### Network architecture and maturation in term-born infants

Overall we find a robust, modular, symmetrical functional organisation of the brain at TEA. Our results confirm and further elucidate the primary-to-higher-order maturational sequence of RSN development.

### Primary networks

We identified five primary RSNs (**Fig. 1A**) which showed adult-like topology from the earliest ages studied (**Fig. 3A**) and no significant change in architecture from 37-43.5 weeks PMA. Primary, unimodal RSNs mature earlier than higher-order networks in the preterm brain (Doria et al., 2010; Liu et al., 2008; Smyser et al., 2010); our finding of an adult-like configuration of primary RSNs at TEA is in agreement with previous studies at this age (Doria et al., 2010; Fransson et al., 2011; Fransson et al., 2009; Fransson et al., 2007; Gao et al., 2014; Gao et al., 2015; Smyser et al., 2010). The precise localisation of sensorimotor networks along the central sulcus is especially striking in our data, even in the youngest infants studied (**Fig. 3A**). Determination of somatotopic maps in primary sensorimotor cortical areas occurs as early as mid-third trimester equivalent age, with similar stimulation response to adults observed by TEA (Allievi et al., 2015; Dall’Orso et al., 2018). We additionally observed a significant increase in core network strength within the somatosensory network from 37-43.5 weeks PMA (**Fig. 4**), possibly reflecting increasing integration of secondary somatosensory cortex at this age (Allievi et al., 2015), and/or increased influence of ex-utero experience on this network. The bilateral insula (**Fig. 1A**) and thalamus (**Supplementary Fig. S1C**) were strongly connected within the medial motor network, consistent with previous studies finding strong thalamocortical connectivity in sensorimotor networks (Smyser et al., 2016; Toulmin et al., 2015).

### Association networks

We identified six RSNs representing higher-order association networks (**Fig. 1C**). Using quantitative (**Fig. 2, Fig. 4**) and qualitative (**Fig. 3**) methods, we found modest expansions in both the spatial extent and core temporal coherence of higher-order association networks from 37-43.5 weeks PMA. To our knowledge this is the first time these changes have been quantified over this brief but developmentally critical period. The heterogeneous timing of functional network development, in which primary networks mature earlier than higher-order association networks, can be related to parallel changes in brain structure (reviewed in (Keunen et al., 2017). Structural connectivity of the cortex begins with thalamic connections to frontal, auditory, visual and somatosensory cortices at 24-32 weeks gestation, while long-range cortico-cortical connections are not established until 33-35 weeks (reviewed in (Dubois et al., 2016; Kostovic and Jovanov-Milosevic, 2006). The same sequence is later repeated in cortical myelination, with the “primordial” sensorimotor and visual cortices histologically more mature at the time of birth (Flechsig, 1901). White matter tracts connecting to these regions, such as the corticospinal tract and optic radiation, are also the first to mature later in infancy (reviewed in (Dubois et al., 2014). The structural and functional ontogeny mirrors the observed behavioural sequence of developmental ‘milestones’ in young children, in which sensorimotor, auditory and visual competencies are acquired before higher-order cognitive functions (Keunen et al., 2017).

The two RSNs showing greatest increase in intrinsic connectivity (core network strength) from 37-43.5 weeks PMA were the posterior parietal network and visual association network (**Fig. 4**). The former encompasses the medial precuneus and posterior cingulate cortices (**Supplementary Fig. S1C**), an area of emerging functional connectivity at TEA (Gao et al., 2009). In adulthood these regions are a prominent component of the DMN, leading some to label infant RSNs encompassing these as DMN precursors (Doria et al., 2010; Fransson et al., 2009; Fransson et al., 2007; Smyser et al., 2010; Smyser et al., 2016). However, the mature DMN also incorporates distinct modules in the anterior cingulate/medial prefrontal cortex, orbitofrontal cortex, lateral temporal cortex and hippocampus (reviewed in (Raichle, 2015)). We observe no temporal involvement in the posterior parietal network, and only sparse frontal involvement, specifically at the right anterior cingulate (**Supplementary Fig. S1C**) and bilateral orbitofrontal cortex (**Supplementary Fig. S1E**). This dominant posterior hub with limited frontoparietal connectivity bears more similarity to the adult DMN under anaesthesia (Amico et al., 2014; Bonhomme et al., 2016). Overall we find support for the concept of fragmented local modules prevailing over long-range integration at this period of development, preceding the emergence of a full analogue of the adult DMN at 6-12 months of age (Gao et al., 2014; Gao et al., 2015; Gao et al., 2009).

The visual association network comprises lateral occipital (**Supplementary Fig. S1D)** and inferotemporal (**Supplementary Fig. S1B**) cortices; regions which contribute to the ventral stream of visual processing, in which simple features coded by primary visual cortex are transformed into higher-level representations of objects, invariant of their size, rotation and position, enabling downstream object recognition and semantic processing (DiCarlo et al., 2012; Goodale and Milner, 1992). It was therefore not surprising to find significant growth in the strength of this network from 37-43.5 weeks PMA (**Fig. 2**, **Fig. 4**), a period in which infants are increasingly exposed to, and able to resolve, objects in the visual field (Dubowitz et al., 1983). Furthermore, after controlling for differences in age, we found increased connectivity within this network of inferotemporal regions including the posterior fusiform gyrus in female infants (**Fig. 5**). The fusiform is sensitive to complex visual stimuli including faces and facial expressions (Li et al., 2019); in the corresponding region of the macaque brain the code determining face cell firing was recently deciphered (Chang and Tsao, 2017). In humans, reduced functional connectivity of the fusiform face area is associated with developmental prosopagnosia (Lohse et al., 2016). The sex difference in functional connectivity we have identified in this region is especially interesting in the context of behavioural data in which female neonates, compared to males, show increased preference for looking at faces (Connellan et al., 2000). A recent study comparing neonates at high familial risk for autism with controls also identified a significant group difference in this region (Ciarrusta et al., 2019). Further investigation of functional connectivity in the ventral stream and social-cognitive development might elucidate mechanisms for sex differences in this domain.

Two RSNs comprised segregated (i.e. non-contiguous) brain regions revealing anatomically meaningful patterns of functional connectivity. The temporoparietal network (**Fig. 1D**) connects a posterior module encompassing the extended Wernicke’s area to a smaller anterior module corresponding to Broca’s area. Integrated structural-functional analysis in adults showed this network is facilitated by the arcuate fasciculus (O’Muircheartaigh and Jbabdi, 2018). The instatement of a putative ‘language network’ in early infancy is supported by stimulus-fMRI showing activation of these regions in response to speech (Dehaene-Lambertz et al., 2002; Dehaene-Lambertz et al., 2006). The frontoparietal network (**Fig. 1B)** connects the frontal, supplementary and parietal eye fields, with close resemblance to the adult dorsal attention network (Vossel et al., 2014). Ocular control relies on widespread white-matter connections between cortical and subcortical regions, the microstructural integrity of which correlates with visual fixation behaviour in the neonate (Stjerna et al., 2015). Striatal projections of the frontal and supplementary eye fields converge upon the caudate nucleus (Parthasarathy et al., 1992); we found a positive association between older PMA at scan and functional connectivity of the caudate nucleus within this frontoparietal network (**Fig. 2**), consistent with active development of the oculomotor corticostriatal system at this age.

### Impact of preterm birth

Preterm birth confers a high risk of neurodevelopmental impairment (Bhutta et al., 2002; Marlow et al., 2005; Saigal and Doyle, 2008) and psychiatric illness in later life (Nosarti et al., 2012). Pervasive deficiencies and delays in structural brain maturation have been identified in preterm infants scanned at TEA, even in those without focal brain injury, including macrostructural differences in tissue volume and gyrification (Ball et al., 2012; Kapellou et al., 2006; Keunen et al., 2012; Shimony et al., 2016) and microstructural alterations in both grey and white matter (Ball et al., 2013b; Bouyssi-Kobar et al., 2018a; Krishnan et al., 2007). The overall structural network architecture appears unchanged, with preservation or even abnormal strengthening of the rich-club organisation of highly connected cortical hubs, at the expense of diminished peripheral connectivity and specific disruptions to thalamocortical, cortical-subcortical and short-distance corticocortical connectivity (Ball et al., 2014; Ball et al., 2013a; Ball et al., 2015; Batalle et al., 2017; Lee et al., 2019).

### Widespread impairment of functional connectivity

Now we show that, similar to structural connectivity, functional connectivity is profoundly affected by preterm birth. We found striking deficiencies in within-network connectivity across the full range of RSNs studied (**Fig. 6**), also replicated as a dose-dependent relationship, such that increased exposure to prematurity (younger GA at birth) was associated with decreased functional connectivity (**Supplementary Fig. S2**). This suggests that although functional connectivity increases across the preterm period (Cao et al., 2016; Doria et al., 2010; Smyser et al., 2010; Smyser et al., 2016; van den Heuvel et al., 2015), it does not reach a normal configuration at TEA. Instead there appears to be an aberrant developmental trajectory, in which connections between brain regions are reconfigured by premature exposure to the extrauterine environment. Graph theoretical approaches have shown global network measures of clustering, integration and modularity at TEA are all reduced in preterm infants compared to full-term controls (Bouyssi-Kobar et al., 2019; Scheinost et al., 2016). Hypothesis-driven seed-based approaches have identified disrupted thalamocortical connectivity (Smyser et al., 2010; Toulmin et al., 2015), consistent with structural disruption of the same (Ball et al., 2015). In our data-driven, whole-brain ICA approach, the main finding was globally reduced within-network functional connectivity. Primary and association RSNs appeared to be similarly affected, in contrast to the findings of Smyser and colleagues, who also employed whole-brain correlation mapping, and found primary RSNs were less affected by prematurity (Smyser et al., 2016). This discrepancy may be due to differences in approach to RSN definition (adult-derived RSNs), network mapping (node-based) and inclusion criteria for the preterm group (< 30 weeks GA at birth). In another study investigating preterm-at-term infants with whole-brain ICA, the method comprised identification of 71 nodes by ICA followed by subject-specific network estimation and selection of discriminatory edges between cases and controls using machine-learning classifiers (Ball et al., 2016). Connections to frontal and basal ganglia nodes were overrepresented among the discriminatory edges, indicating altered connectivity in preterm infants. Taken together, these different approaches provide complementary demonstrations of spatially widespread impaired RSN coherence in the preterm-at-term brain.

### Modulation of parieto-motor connectivity

In the context of brain-wide deficiencies in functional connectivity in preterm-at-term infants, it was notable that there was also increased functional connectivity of the bilateral superior parietal lobule (Brodmann area 5) within the lateral motor network, both when prematurity was evaluated as a group effect (**Fig. 6**) and as a continuous variable (**Supplementary Fig. S2**). The lateral motor network corresponds approximately to the primary somatotopic regions serving the upper limb, hand and face (**Fig. 1D**). Ex-utero experience during the preterm period strongly influences the development of sensorimotor networks: bilateral functional responses in the perirolandic cortices to stimulation of the wrist increases with postnatal age, even after controlling for GA at birth (Allievi et al., 2015). Interestingly, connectivity with superior parietal regions appears to occur as a feature of normal development in the lateral motor network in older term-born infants (**Fig. 3A**). Area 5 comprises the somatosensory association cortex which integrates visual and somatosensory inputs to encode limb configuration in space, enabling coordinated movements within the immediate environment (Graziano et al., 2000; Mountcastle et al., 1975). It is intuitive that connectivity of area 5 with lateral motor cortex could be highly dependent upon ex-utero experience, given the natural constraints upon limb movement and visuomotor integration in utero. We propose therefore that the experience of premature exposure to the extrauterine environment modulates the normal development of parieto-motor connectivity, leading to an abnormal increase in connectivity at TEA.

Previous studies have identified increased functional connectivity of certain primary cortical regions in preterm-at-term infants compared to controls, specifically the lateral postcentral gyrus with the thalamus (Toulmin et al., 2015) and regional connectivity within occipital/visual networks (Bouyssi-Kobar et al., 2018a). This may occur at the expense of connectivity in other brain areas, and can persist into later life; analysis of language networks in preterm children scanned at 12 years of age showed increased connectivity with primary sensorimotor areas, but reduced connectivity with higher-order frontal areas (Schafer et al., 2009). Relatively conserved topology of core structural networks has been reported in preterm-born babies (Batalle et al., 2017), persisting into later childhood and adulthood (Fischi-Gomez et al., 2016; Karolis et al., 2016). Disruption of the normal balance of sensorimotor development may have persisting effects on later motor and cognitive development. In the mature brain, the superior parietal lobule supports not only the smooth execution of motor plans (Simon et al., 2002) but also more abstract visuospatial functions such as mental rotation (Gogos et al., 2010). The aberrant parietal connectivity we have identified at TEA could therefore be a prelude to specific difficulties occurring with high prevalence in preterm children, such as developmental coordination disorder (Caravale et al., 2019; Davis et al., 2007; Dewey et al., 2019; Kashiwagi et al., 2009; Wilson et al., 2017), inattention and intellectual impairment (reviewed in (Rogers et al., 2018). Long-term follow up of the study population at school age will be required to confirm this hypothesis.

### Limitations

The customised neonatal imaging system for the dHCP includes a close-fitting head coil sized specifically for the neonatal head, thus providing exceptional signal-to-noise at the cortical surface (Hughes et al., 2017). This bias towards surface-proximate sources is compounded by the use of highly accelerated multiband EPI (Fitzgibbon et al., 2019). As such, this has likely resulted in greater sensitivity to detect correlated signal fluctuations in the cerebral cortex compared to deeper sources such as the thalamus, basal ganglia and cerebellum. This may explain the relatively sparse involvement of subcortical regions in the identified RSNs (**Fig. 1A, 1C**). Thalamocortical and cerebellar functional connectivity may be better appreciated with seed-based methods (Herzmann et al., 2018; Toulmin et al., 2015). We also noted sparse involvement of inferior frontotemporal regions, even at *Z* > 1 (**Supplementary Fig. S1B**). The dHCP functional pipeline includes advanced distortion-correction techniques (Fitzgibbon et al., 2019), but some signal loss related to air/tissue and bone/tissue interfaces in this vicinity cannot be fully excluded. However, this may also reflect biological reality in these brain regions, which are the least myelinated at birth (Flechsig, 1901) and so may be the least able to participate in long-range phase-synchronous activity. In this study we used a dense sampling strategy at TEA to infer longitudinal change in RSNs, but each infant was scanned on only one occasion. GA at birth and PMA at scan were strongly correlated within the term-born group, which complicates the interpretation of these longitudinal analyses. Furthermore, as some potentially relevant neonatal characteristics such as intracranial volume and postnatal days of life, are intrinsically associated to some of our variables of interest (i.e., postmenstrual age at scan, sex, gestational age at birth), it is difficult to disentangle their relative contributions to our results.

The optimised fMRI dHCP pipeline includes multiple steps to control for motion and physiological confounds, thus minimising data loss. However, while well-fed babies tend to fall asleep during the scan, subject motion is inherently correlated with the arousal and sleep state of the baby, which may have an effect in the reconstructed RSNs (Horovitz et al., 2009). While our stringent control for high motion during the scan will minimise the potential effect of subject differences in arousal and sleep state, the specific measure that should be used as a surrogate to model arousal state is unclear. Future studies using simultaneous EEG-fMRI could help to better understand the effect of different sleep states on RSNs. Differences in arousal in the scanner between infants and adults should also be considered when comparing RSN topology between these groups (Mitra et al., 2017). Our use of infants scanned at 43.5-44.5 PMA to define the group-ICA components may have missed some sources of structured noise occurring predominantly at younger ages, such as CSF signal in the cavum septum pellucidum. More fundamentally, the extent to which BOLD signal might be confounded by cerebrovascular factors differing between preterm- and term-born infants (Bouyssi-Kobar et al., 2018b) remains open to debate. Some of the important temporal dynamics in functional networks may be missed by rs-fMRI, which predominantly identifies activity at < 0.1Hz (Cordes et al., 2001). Complementary approaches such as EEG may help to address this (Arichi et al., 2017; Mehrkanoon, 2019).

## Conclusion

Brain development occurs in a pre-programmed and spatially heterogeneous progression, modulated by environmental influence. As such, we observed different trajectories for different neural systems, obeying a generally primary-to-higher-order sequence of maturation. At TEA we find already instated a complete set of adult-analogous unimodal RSNs corresponding to primary sensorimotor, visual and auditory cortices, with relatively little change from 37-43.5 weeks PMA. In contrast, association RSNs appear fragmented and incomplete compared to the adult repertoire, and are undergoing active maturation at this time. Connectivity within the visual association network in particular is highly associated with age, likely as a result of postnatal environmental experience, but also modified by the sex of the infant. Preterm birth is associated with profoundly reduced functional connectivity across all RSNs, but also with augmentation of parieto-motor connectivity, with possible implications for understanding certain neurocognitive sequelae of prematurity. In future we may be able to positively modulate RSN development in prematurity via targeted environmental manipulations (Lordier et al., 2019). Preterm birth is best conceptualised as a developmental perturbation which reconfigures, rather than simply diminishes, the organisation of functional brain networks.

## Figure Legends

**Supplementary Figure S1.** Spontaneous BOLD activity patterns (RSNs) derived from group ICA in 24 term-born infants scanned at 43.5-44.5 weeks PMA. RSN are expressed as a functional parcellation of the brain using a ‘winner-takes-all’ approach based on the RSNs from group ICA. RSNs were spatially smoothed and thresholded at Z > 1 prior to determination of the ‘winning’ RSN at each voxel. The resulting volume was projected to the midthickness cortical surface using enclosed (nearest neighbour) volume-to-surface mapping, here displayed on the pial surface of an individual subject scanned at 42 weeks PMA and viewed from the superior (A), inferior (B), medial (C), lateral (D), anterior (D) and posterior (E) aspects.

**Supplementary Figure S2.** Effect of gestational age at birth in functional connectivity. Association of functional connectivity and gestational age at birth (GA) in term- and preterm-born infants scanned at 37-43.5 weeks PMA. Coloured t-statistic maps thresholded at p < 0.025 (FWE corrected) show connectivity in brain regions negatively (blue) or positively (red-yellow) associated with gestational age at birth. Example sagittal, coronal, and axial slices for meaningful spatial patterns within each RSN are shown, overlaid on a T1 structural template and displayed in radiological convention. White lines represent the outlines of the group-ICA RSNs, thresholded at Z > 3

