## Supplemental Figures for "The Developing Human Connectome Project: typical and disrupted perinatal functional connectivity"

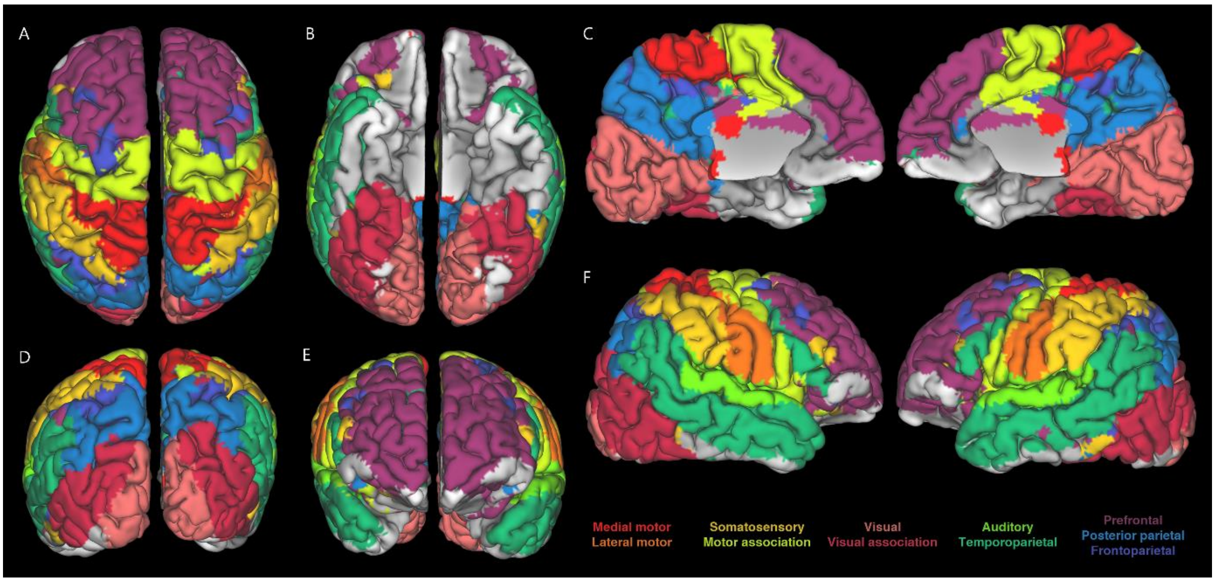


**Figure S1.** *Spontaneous BOLD activity patterns (RSNs) derived from group ICA in 24 term-born infants scanned at 43.5-44.5 weeks PMA. RSN are expressed as a functional parcellation of the brain using a 'winner-takes-all' approach based on the RSNs from group ICA. RSNs were spatially smoothed and thresholded at Z > 1 prior to determination of the 'winning' RSN at each voxel. The resulting volume was projected to the midthickness cortical surface using enclosed (nearest neighbour) volume-to-surface mapping, here displayed on the pial surface of an individual subject scanned at 42 weeks PMA and viewed from the superior (A), inferior (B), medial (C), lateral (D), anterior (D) and posterior (E) aspects.*

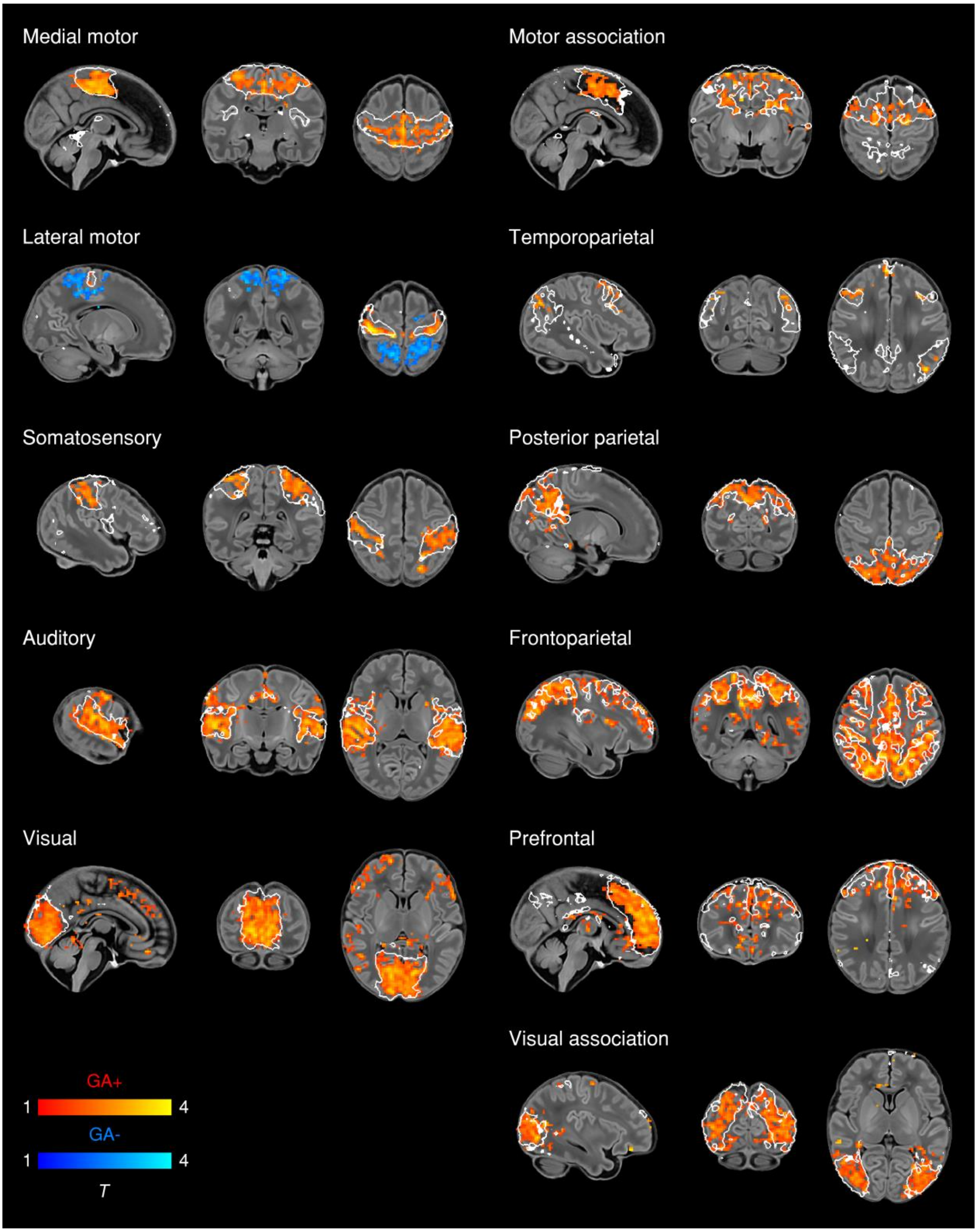


**Figure S2.** ***Effect of gestational age at birth in functional connectivity.*** *Association of functional connectivity and gestational age at birth (GA) in term- and preterm-born infants scanned at 37-43.5 weeks PMA. Coloured t-statistic maps thresholded at $p < 0.025$ (FWE corrected) show connectivity in brain regions negatively (blue) or positively (red-yellow) associated with gestational age at birth. Example sagittal, coronal, and axial slices for meaningful spatial patterns within each RSN are shown, overlaid on a T1 structural template and displayed in radiological convention. White lines represent the outlines of the group-ICA RSNs, thresholded at $Z > 3$.*
